# Sexual conflict mitigation via sex-specific trait architecture

**DOI:** 10.1101/2022.11.30.518524

**Authors:** Simona Kralj-Fišer, Matjaž Kuntner, Paul Vincent Debes

## Abstract

Sexual dimorphism — the sex-specific trait expression — may emerge when selection favours different optima for the same trait between sexes, i.e., under antagonistic selection. Intra-locus sexual conflict exists when the sexually dimorphic trait under antagonistic selection is based on genes shared between sexes. A common assumption for sexual-size dimorphism (SSD) is that its presence indicates resolved sexual conflict, but how current sex-specific evolution proceeds under sexual dimorphism remains enigmatic. We investigated whether a sex-specific architecture of adult body size explains sexual conflict resolution under extreme SSD in the African hermit spider, *Nephilingis cruentata*, where adult female body size greatly exceeds that of males. Specifically, we estimated the sex-specific importance of genetic and maternal effects on adult body size among individuals that we laboratory-reared for up to eight generations. Quantitative genetic model estimates indicated that size variation in females is to a larger extent explained by direct genetic effects than by maternal effects, but in males to a larger extent by maternal than by genetic effects. We conclude that this sex-specific body-size architecture enables body-size evolution to proceed much more independently than under a common architecture to both sexes, thereby mitigating sexual conflict under SSD.

## Introduction

Sexual dimorphism, the between-sex trait difference in a trait, exists in many animals. How evolution of existing sexually dimorphic traits may proceed has long been a mystery to researchers [1]. Specifically, selection may favour sex-specific optima in the same trait, generating antagonistic selection between the sexes, and this trait may be determined by shared genes between the sexes, defining intra-locus sexual conflict [2-4]. Sexual conflict and its mitigation, or resolution, not only play an important role in the evolutionary emergence of sexual dimorphism and have far-reaching effects on genomic organization and speciation, but also on processes that act on medium to short scales, such as population dynamics or extirpation [5-11]. It may thus not be surprising that elucidating how sex-specific evolution of sexually dimorphic traits proceeds has been subject to much past and current research [3, 12-16].

Sexual dimorphism in size, termed sexual-size dimorphism (SSD), may have resulted from sex differences in the optimal body size relating to either parental investment or mating success [1, 14, 17]. Specifically, anisogamy – the differences between sex-specific gametes – often requires higher energetic investment by females to produce eggs (or offspring) than sperm cells produced by males [17, 18]. SSD with females as the larger sex (female-biased SSD) may then have emerged in systems in which female, but not male, body size affects offspring size and number [1, 19-21]. However, the genetic and molecular mechanisms that allow sex-specific evolution of sexually dimorphic traits remain largely unknown, although it is often assumed that the presence of SSD implies at least partly resolved sexual conflict [3, 14, 20]. A common assumption is that sexual-conflict resolution involves a decoupling of the genetic architecture between the sexes [11, 22].

Exactly how decoupling of the genetic architecture between sexes proceeds to allow for an independent evolution of the sexes is subject to current research. Theoretically, sexual conflict can be resolved by mechanisms leaving distinct signatures that can be detected using quantitative genetic methods. Specifically, a resolution may lead to detecting heterogeneous direct genetic variances between sexes or a low between-sex genetic correlation [6]. However, the between-sex genetic correlation may often, but not always, predict the degree of sexual dimorphism [16, 23] making it worthwhile to consider resolving mechanisms involving effect levels other than the direct genetic, such as the maternal effect level [24, 25]. Maternal effects, i.e., causal influences of the maternal phenotype on the offspring phenotype other than that of her directly transmitted genetic variants, may vary with maternal environment or maternal genetics [26-29]. Whereas maternal *environmental* effects on offspring are controlled by the maternal environment on a maternally expressed trait, maternal *genetic* effects on offspring are controlled by direct genetic effects on a maternally expressed trait. Only the latter effects are heritable. Sex-specific maternal effects, although empirically associated with sexual dimorphism in only a few cases [30-32], have long been considered theoretically in selection for sexual dimorphism [33] and more recently in the resolution of sexual conflict [24, 25]. Importantly, if variation for body size between the sexes underlies different relative contributions of maternal and direct genetic effects, this would enable resolving, or at least mitigating, sexual conflict over body size and thus maintaining SSD while allowing for sex-specific evolution of body size.

Here, we examined sex-specific adult body size variation in the African hermit spider, *Nephilingis cruentata*, which expresses an extremely female-biased SSD [34, 35]. We reared spiders for up to eight generations under standardized laboratory conditions, measured 2,540 pedigreed individuals, and tested, using quantitative genetic methods, whether the relative contributions by direct genetic and maternal effects to adult body mass variation differed between sexes. Our results suggest that variation in adult body mass is explained to a larger extent by direct genetic effects in females and to a larger extent by maternal effects in males, whereby direct genetic effects may play a minor role on size variation of males. Our results support the presence of a relatively straightforward mechanism that mitigates intra-locus sexual conflict and allows for a less constrained sex-specific evolution of adult body size than under a common body size architecture.

## Results

### Adult body size varies with season

We first evaluated whether the day of the year when spiderlings hatched affected their adult body size, because adult body size of many spiders is influenced by seasonal environmental factors [35-37]. We were concerned that unaccounted seasonal effects across the six years of rearing, but common to concurrently hatching siblings, might be confounded with (other) maternal effects. The results of a mixed model accounting for relatedness via the inverse of the additive genetic relatedness matrix and maternal effects via mother identification indicated that adult body mass indeed associated with hatching season in both sexes (**Figure 1, Table 1**). Specifically, individuals hatched during summer were larger than those hatched during winter, whereas individuals hatched during spring and autumn expressed intermediate body sizes. Further, average adult body size increased with decreasing daylength when hatched in summer and decreased with increasing daylength when hatched in winter.

**Table 1.** ANOVA table for fixed effects in the mixed model for adult body mass of female and male spiders.

| Term | DF | DDF | F | P |
| --- | --- | --- | --- | --- |
| *Sex* | 1 | 12.5 | 9818 | < 0.001 |
| *Period* | 1 | 106.1 | 1 | 0.424 |
| *Sex-by-Period* | 1 | 145.7 | 4 | 0.057 |
| *Maternal food* | 1 | 291.2 | 1 | 0.364 |
| *Sex-by-Maternal food* | 1 | 288.3 | 1 | 0.357 |
| *Date*, polynomial order 1 | 1 | 240.8 | 82 | < 0.001 |
| *Date*, polynomial order 2 | 1 | 260.8 | 0 | 0.560 |
| *Date*, polynomial order 3 | 1 | 262.6 | 32 | < 0.001 |
| *Sex-by-Date*, polynomial order 1 | 1 | 200.5 | 1 | 0.475 |
| *Sex-by-Date*, polynomial order 2 | 1 | 233.8 | 12 | < 0.001 |
| *Sex-by-Date*, polynomial order 3 | 1 | 205.4 | 0 | 0.777 |

Terms: *sex* (female, male), *Period* relative to introducing the maternal food treatment (before, after), *Maternal food* after reaching adulthood (high, low), *Date* as day of year when hatched (continuous: 1-366); DF: degrees of freedom; DDF: denominator degrees of freedom. The *Period* term was fitted to enable a direct testing of the *Maternal food* level high-low contrast and its interaction with *Sex*.

**Figure 1.**
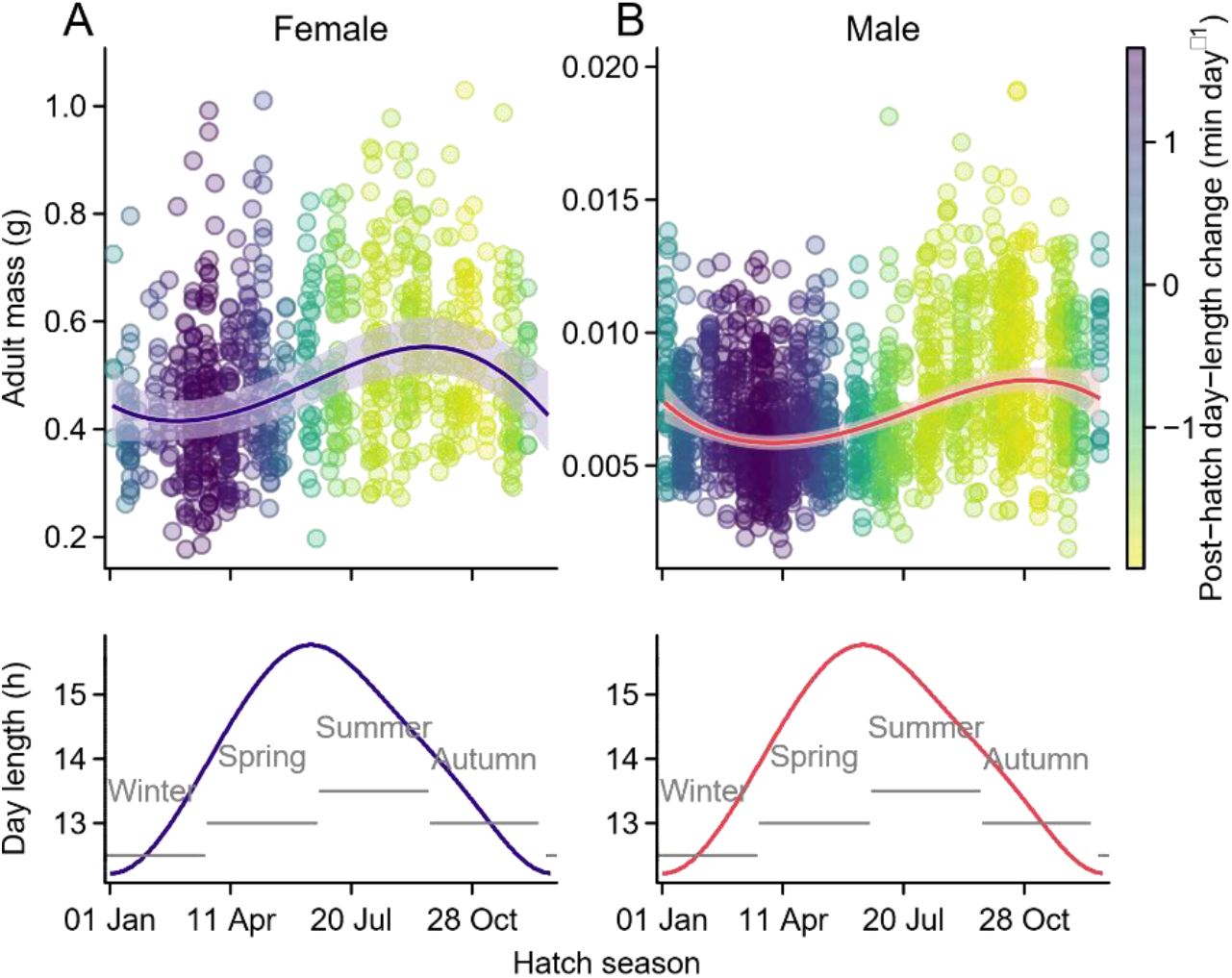
Seasonal change of adult body mass. Model-predicted trends of adult body mass in female (**A**, *n* = 789) and male (**B**, *n* = 1,751) hermit spiders across seasons (when hatched). Lines with 95% confidence bands represent the 3^rd^ order polynomial model fit for adult mass across hatching day of year in females (purple) or males (red). Points represent individual measurements with colour indicating the average change in daylength during the first seven days after hatching. Effective daylength per day and seasons are shown in the bottom panels.

Using the same mixed model, we tested whether feeding prospective mothers one fly (low food treatment) or three flies (high food treatment) twice per week after mating affected adult body size of their daughters or sons, thereby testing for sex-specific maternal *environmental* effects related to maternal food amount. We did not detect convincing evidence that the maternal food amount affected body size of offspring of either sex. Specifically, both the main and interaction terms of the maternal food treatment with sex were non-significant (**Table 1**). Daughters from high-food mothers were estimated to be only 1.07 times (95% confidence interval, 95% CI: 0.97-1.18 times) larger compared with daughters from low-food mothers, and this difference was non-significant (*t*_291.2_ = 1.30, *P* = 0.197). Sons from high-food mothers were estimated to be of very similar size to sons from low-food mothers, specifically just 1.01 times (95% CI: 0.93-1.10 times) larger, which was also non-significant (*t*_291.2_ = 0.21, *P* = 0.834).

### Body size architecture differs between sexes

Controlled for sex, hatching season, and maternal food treatments, we observed opposite importance between sexes for direct genetic vs. maternal proportional contribution-estimates to variation for adult body mass (heritability, ĥ^2^, and 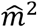, respectively; **Figure 2**; (co)variances in **Table 2**).

**Table 2.** Sex-specific variance estimates of adult body mass for either females (F) or males (M), and between-sex covariance estimates, for maternal (*dam*), direct genetic (*animal*), or residual effects by either REML or parametric bootstraps.

| Term | REML |  | Boot |  | 2.50% | 97.50% |
| --- | --- | --- | --- | --- | --- | --- |
|  | Estimate | SE | Estimate | SE |  |  |
| *dam*, $\hat{\sigma}_F^2$ | 0.0104 | 0.0041 | 0.0102 | 0.0038 | 0.0032 | 0.0182 |
| *dam*, $\widehat{cov}_{F,M}$ | 0.0080 | 0.0031 | 0.0077 | 0.0029 | 0.0022 | 0.0134 |
| *dam*, $\hat{\sigma}_M^2$ | 0.0246 | 0.0042 | 0.0234 | 0.0037 | 0.0163 | 0.0308 |
| *animal*, $\hat{\sigma}_F^2$ | 0.0213 | 0.0084 | 0.0224 | 0.0087 | 0.0066 | 0.0411 |
| *animal*, $\widehat{cov}_{F,M}$ | -0.0002 | 0.0048 | 0.0005 | 0.0045 | -0.0079 | 0.0099 |
| *animal*, $\hat{\sigma}_M^2$ | 0.0014 | 0.0054 | 0.0043 | 0.0039 | 0.0000 | 0.014 |
| *residual*, $\hat{\sigma}_F^2$ | 0.0514 | 0.0054 | 0.0510 | 0.0055 | 0.0398 | 0.0615 |
| *residual*, $\hat{\sigma}_M^2$ | 0.0720 | 0.0037 | 0.0706 | 0.0033 | 0.0638 | 0.0768 |

**Figure 2.**
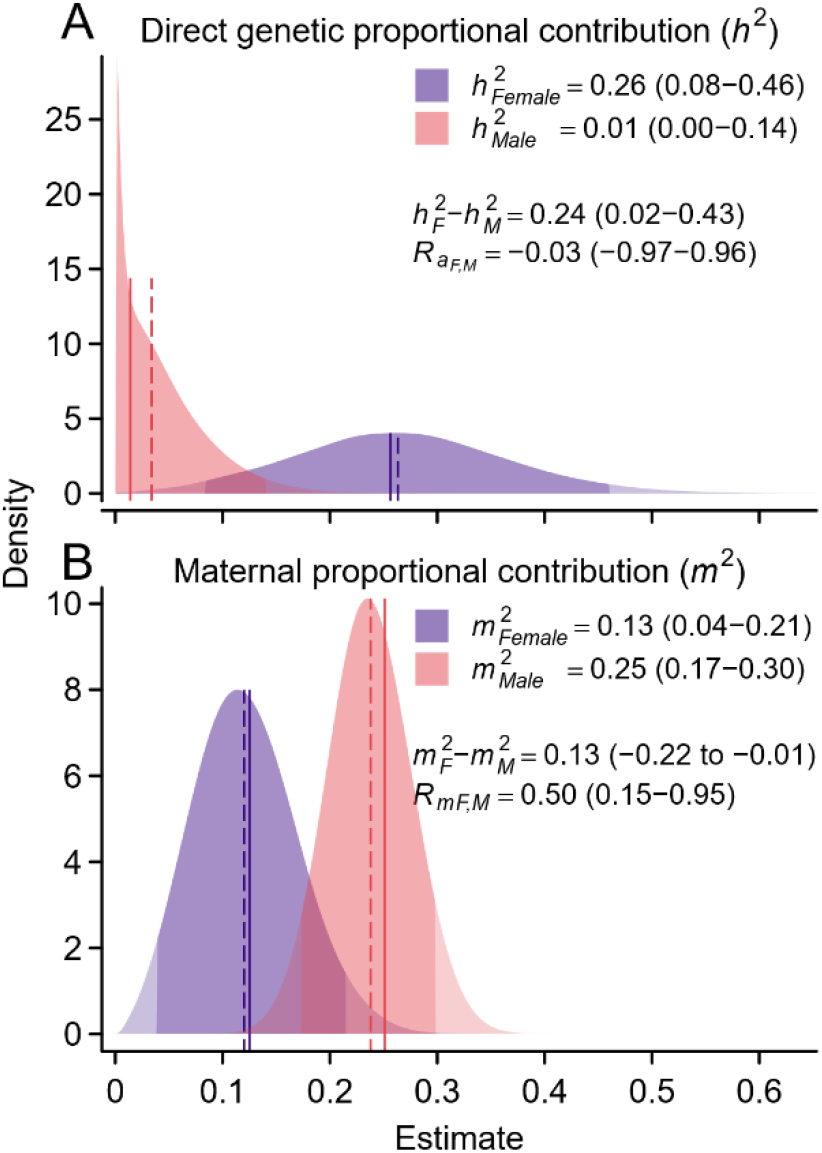
Sex-specific body size architecture. Sex-specific estimates of proportional contribution to the phenotypic variance of adult body mass by direct genetic effects (*h*^*2*^; **A**) and maternal effects (*m*^*2*^; **B**) in female (purple; *F*) and male (red; *M*) African hermit spiders. Means were estimated by REML, whereas the 95% confidence intervals for each distribution, indicated by a stronger colour saturation, were estimated across 10,000 parametric bootstrap replicates. REML means and bootstrap medians are indicated by vertical solid and dashed lines, respectively.

Specifically, in females (*F*), direct genetic effects made up 26% of the phenotypic variance, but maternal effects made up only 13%, and the lower confidence interval for both estimates were well away from zero. In contrast, in males (*M*), direct genetic effects made up only 1% of the phenotypic variance, whereas the maternal effect variance made up 25%, and the lower confidence interval of the former but not the latter approached zero. This opposite importance between sexes for relative amounts contributed by genetic (*a*) vs. maternal (*m*) effects on variation of body size phenotype expression was supported by the 95% confidence intervals for the between-sex contrasts of heritability and the maternal proportional variance contribution that both excluded zero (**Figure 2;** see tests on variance differences below). Under sex-specific optima of the same trait, sexual conflict may be detected by a high and positive genetic correlation between the sexes, as it can constrains the sex-specific evolution of a trait by inducing correlated selection responses of the two sexes [6, 38, 39]. The between-sex correlation estimates for the direct additive genetic correlation 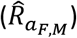 showed, albeit estimated as close to zero, a wide confidence interval spanning both negative and positive values (**Figure 2**). This, however, is not unexpected when male direct genetic effect estimates have a large uncertainty relative to their estimates (i.e., under a low genetic variance; **Table 2**), so that their ranking and thus correlation with the female effects is uncertain. However, the between-sex correlation estimate for maternal effects 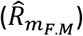 and its 95% confidence interval were positive, indicating that maternal effects are – despite showing a differential relative importance – shared to some extent between sexes.

Comparisons of estimates for the proportional contribution to the phenotypic variance, such as ĥ^2^, and 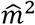, may not fully reflect the differences in evolvability [40], but instead for log transformed trait data the differences in genetic variance estimates may be preferred [41]. Therefore, we tested the hypotheses of sex differences in variance estimates using likelihood ratio tests between the model with sex-specific variances and each of three nested models in which we constrained genetic, maternal, or residual variance to be the same for the sexes. We found the model fitting different genetic variances between sexes (15.5 times larger for females) to be better than the model fitting a genetic variance constrained to be the same for the sexes 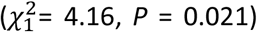. This leads to the expectation that a proportional response (i.e., considering the different absolute body sizes between sexes) to a selection gradient with identical values for the sexes on the log scale would incur a larger response for females than males. In addition, we found the model with sex-specific maternal variances (2.4 times for males) and residual variances (1.4 times larger for males) to fit better than models fitting each of these variances constrained to be the same for the sexes (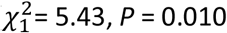 and 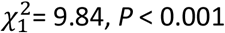, respectively).

We also tested how much the included fixed effects (hatching season, maternal food treatment) affected the proportional contribution and variance estimates. Not controlled for hatching season, heritability estimates decreased slightly in females and increased slightly in males (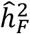: changed from 26% to 18%; 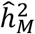: 1% to 3%), and the uncertain between-sex genetic correlation estimate decreased considerably (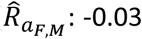 to -0.88), whereby the latter may have been a statistical consequence of the abovementioned low male effect variance. In contrast, the proportional contribution of maternal variance increased – as expected – in both sexes (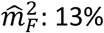 to 25%; 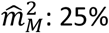 to 32%). The changes in proportional contributions were caused by slightly lower and higher direct genetic variance estimates in females and males, respectively, and noticeable higher maternal variance estimates in both sexes when not accounting for hatching season (electronic supplementary material, **Appendix 1 - figure 4**). Along with increased maternal effect variance, the between-sex correlation for maternal effects increased (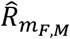: from 0.50 to 0.69). We thus confirmed that non-controlled hatching-season effects manifest, statistically, as common environmental effects that are correlated between sexes (detected in simultaneously hatching siblings as maternal environmental effects) and contribute about 7-12% to the phenotypic variance. Not controlled for the maternal food treatment (i.e., pooling high and low treatments), the proportional contribution of direct genetic and maternal effects to the phenotypic variance mirrored estimates obtained when the treatments were controlled for, except for a somewhat lower female heritability (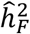: from 26% to 24%) caused by a somewhat lower direct genetic variance in females (electronic supplementary material, **Appendix 1 - figure 4**). Likewise, the between-sex correlations for genetic and maternal effects were comparable to the estimates by the full model (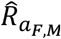: from -0.03 to -0.07; 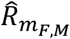 :0.50 to 0.48). We thereby confirmed the results obtained when testing maternal food treatments as fixed effects and gathered evidence that the maternal food treatments may have had little (or no) effects on the maternal variance estimates of either sex.

## Discussion

We here inferred that adult body size variation in a non-model species with extreme sexual-size dimorphism (SSD) is explained to a larger extent by direct genetic effects in the larger females and to a larger extent by maternal effects in the ∼75 times smaller males for which direct genetic effects appeared to play a minor or no role on size variation. These results on a spider species support the hypothesis that sexual conflict can be resolved, and SSD maintained while allowing for current sex-specific evolution through sex-specific trait architecture. Simply put, the documented architecture of adult body size implies that size variation in daughters underlies to a large share by the directly inherited alleles from both parents. In contrast, adult body size variation in sons appears to underlie little on the directly inherited alleles from either parents that are expressed in the offspring. Instead, adult body size architecture of males appears to be influenced by an unknown trait expressed in their mother (or of common environmental effects to siblings from the same egg sac), and this maternal trait may or may not have a direct genetic basis in the mother. Regardless of whether the maternal effect has a direct genetic basis in the mother, to the offspring it acts as an environmental effect independent of the genes inherited by either parent [29]. The genes inherited by both parents may have a low importance on adult male size variation, as we estimated both a low heritability and a low log-scale direct genetic variance. This sex-specific architecture of adult body size, altogether, implies a resolved intra-locus conflict that allows size evolution to proceed at the direct genetic level in females, with minor consequences on male size. These results on quantitative genetic parameters provide evidence how intra-locus sexual conflict can be mitigated or even resolved by sex-specific architecture and thus explain how evolution toward sex-specific optima of the same trait is possible while maintaining sexual dimorphism.

A sex-specific trait architecture is one of several mechanisms circumventing the genetic constraints imposed when a single sexually dimorphic trait underlies shared genes between sexes, i.e., to resolve intra-locus sexual conflict. Proposed mechanisms also comprise effects beyond direct genetic inheritance, including maternal effects [24, 30, 32, 42]. However, empirical studies have remained scarce and provided only limited evidence for sex-bias in maternal effects on sexually dimorphic traits [30, 43-46]. In the current study, maternal effects explain 13% of the phenotypic variance (i.e., *m*^*2*^) of adult body size in females. In contrast, in males the estimate was 25% and likelihood ratio tests indicated that the maternal variance estimates (adjusted for size differences) were larger in males than females. In addition to this relatively small sex-bias for variation in maternal effects, we detected a substantial sex-bias for variation in direct additive genetic effects. Variation in direct genetic effects explained 26% of the phenotypic variance (i.e., *h*^*2*^) in females, which was considerably higher than the estimated 1% in males. A likelihood ratio test indicated here that the log-scale genetic variance estimate was larger in females than males, suggesting a higher evolvability by direct genetic effects in females than males. Together, these empirical results support the idea that sex-specific evolution under extreme SSD is possible through differences in trait architecture between the sexes. In our case, the architecture of adult body size involves direct genetic effects predominantly in females, and maternal effects in both sexes, whereby the latter appeared less important in females than in males. Nonetheless, the results also suggested the presence of a positive between-sex maternal correlation, which we estimated with large uncertainty. It therefore remains unclear whether a sex-specific evolution via maternal effects – if these are maternal genetic effects – is constraint. Regardless of such a possible genetic constraint at the maternal level, the direct genetic effects on body size remain largely restricted to females.

A major question emerging from the results is whether the estimated maternal contribution to adult body size is governed by environmental, genetic, or both effects. Using model selection, we concluded maternal *environmental* effects to fit the data slightly better than maternal *genetic* effects (electronic supplementary material, Model selection), but using data simulations we were unable to fully disentangle these effects from each other (electronic supplementary material, Data simulation, **Appendix 1 - figure 3**). However, the type of maternal effect underlying a trait matters regarding the mechanisms controlling its evolution [26, 29, 46, 47] and may, or may not, encompass the abovementioned constraints on sex-specific evolution via shared genes. Specifically, under maternal environmental determination, the maternal contribution to the adult body size variation of her offspring depends on the environmental conditions experienced by her, which is usually assumed to be independent of the direct genetic determination of her own body size. In contrast, under maternal genetic determination, the maternal contribution to the adult body size variation of offspring depends on allelic variants inherited by both of the parents of the mother (i.e., the grand-parents of an individual) and follows a predictable but by one generation lagged pattern of inheritance and thus response to selection [29]. Perhaps more important for resolving sexual conflict under this scenario, sex-specific maternal genetic and direct genetic effects may be subject to similar mechanisms that constrain sex-specific evolution under a direct genetic architecture for both sexes. In detail, if adult body size is determined by direct genetic effects in females (*a*_*F*_) and by maternal genetic effects in males (*mg*_*M*_), the between-sex correlation of these effects 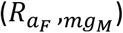 may still point towards an intra-locus sexual conflict because it may indicate shared genes between sexes at the direct genetic and maternal genetic levels [26, 48]. An example would be when the maternal adult body size (controlled by direct genetic effects) affects the maternal genetic effects on adult body size of her sons (controlled by maternal genetic effects). However, when we fitted a more complex (but less supported) model that estimated this correlation (electronic supplementary material, **Appendix 1 - figures 1, 2**), it was estimated to be close to and not different from zero 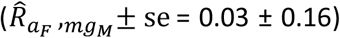.

Assuming a maternal genetic rather than environmental contribution to male adult body size variation, this low or zero between-sex correlation indicates that different gene sets expressed in different generations (mothers vs. offspring) are the major genetic determinants of sex-specific adult body size and that intra-locus sexual conflicts are thus probably largely resolved.

Our results also suggest that sex chromosomes, which determine sex in spiders and have long been thought to play important roles in sexual conflict [15, 49], may here not be very strong candidates for explaining the differences in adult body size architecture between sexes. In the studied spider species, the X_1_X_2_0 sex-chromosome system prevails [50], which is the most common system in spiders [51]. Under this system, sons inherit one chromosome pair from only the mother (i.e., X_1_X_2_), but daughters inherit one chromosome pair each from both the mother and the father (i.e., X_1_X_1_X_2_X_2_). Thus, recombination is possible in heterozygotic females but not in hemizygotic males (no recombination is assumed to occur between X_1_ and X_2_). According to this pattern of inheritance and recombination, the non-recombined sex chromosome pair passed on by the father to only daughters may be expected to leave quantitative genetic signatures of female-limited *paternal* genetic effects. Likewise, the recombined sex chromosome pair passed on by the mother to daughters *and* sons may be expected to leave signatures of similar direct genetic effects that are correlated between sexes. Whereas the first expectation is difficult to test with our data (like for *maternal* genetic effects), at least the latter expectation is inconsistent with the main sex differences in trait architecture inferred here.

To more easily predict evolution of male adult body size under influence of maternal effects, the actual female trait underlying the maternal effects may be identified. The female trait associated with the maternal effects on male (and to some extent female) adult body size in this study remains unknown, did not appear to relate to female food amount after reaching adulthood, but may relate to other known maternal traits that affect offspring size, such as variation in egg quality or size, or amount of egg-deposited RNA or hormones [26, 52]. Regardless of what the unknown maternal trait is, another important question is why adult body size of daughters appears less affected by them. A relatively simple mechanism may relate to the sex-specific developmental durations. In detail, we estimated that females take more than twice as long as males to reach adulthood, namely 219 days in females vs. 90 days in males. Because importance of maternal effects often declines during ontogeny [45, 53], the contribution of maternal effects to adult body size may be expected to be greater in sons than daughters as a simple consequence of the shorter developmental duration to reach the same developmental stage [33, 35].

For spiders, molecular developmental aspects in the control of sexual dimorphism remain largely unknown but have been suggested as promising candidates to provide results that will enrich our understanding of the underlying mechanisms [54]. However, even though an influence of sex-specific developmental duration on general SSD prevails in insects [55], effects of the differences in developmental duration on differences in trait architecture, as may exist here (direct genetic vs. maternal), do appear to have rarely been linked conceptually [35]. Thus, our results also suggest that adding aspects of sex-specific direct genetic vs. maternal effects to studies of the molecular developmental control of SSD may pave future research avenues to an understanding of the molecular mechanisms that enable sex-specific evolution of sexually dimorphic traits.

## Materials and methods

### Study population, mating design, rearing and maternal food treatment

The studied population of the African hermit spider (*N. cruentata*), a species of IUCN least concern [56], has been maintained at the Institute of Biology ZRC SAZU, Slovenia since 2015. It was founded by 23 wild females collected either already gravid in 2015 in iSimangaliso Wetland Park and Ndumo Reserve, South Africa (permit number OP 552/2015 from Ezemvelo KZN Wildlife), or as virgins in 2018, and one virgin male collected in 2018 in iSimangaliso (continuous permit number OP 3031/2020). In *Nephilingis*, both sexes possess paired genitalia and during copulation the used male palp (genitalia) breaks off within the female’s genital opening, impeding re-mating with the used genitalia [57], and limiting the possible individual copulations to two [37]. Although females may practice sexual cannibalism, it is common for a male to guard a subadult female against suitors prior to maturity and after copulation. Mating of both sexes is thus usually limited to one partner (monogamy) [37, 57]. For gravid wild females, we therefore assumed single male partners. In the laboratory, we mated spiders randomly but avoided full-or half-sib matings. Mating in *N. cruentata* usually involves individuals hatched at different times (of different age) because of a much shorter developmental time, and thus generation time, of males than females [35]. We mated all females, except one, with one male each, and 43 males successfully with two females each, and all others with one female each. The final pedigree spans eight generations, encompasses 318 mothers and 273 fathers (including unknown wild ‘phantom’ partners of the gravid wild females) and contains altogether 2,768 entries. Using the R-package purgeR [58], we estimated the pedigree-based effective population sizes (*N*_*e*_) and average inbreeding coefficient 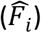 across the last two generations (generations seven and eight) as *N*_*e*_= 62.5 and *F*_*i*_ = 0.048, respectively.

For pairwise mating, we placed an adult female in a poly(methyl methacrylate) frame (35 × 35 × 12 cm) to build a web up to seven days before we added a male using a paint-brush. Because *Nephilingis* males generally mate opportunistically and approach females when disturbed, we placed two to three blow flies (*Lucilia sericata*) on the web for disturbance about 15 minutes after trial commenced. We concluded mating success within 60 minutes, when we placed the female back in her holding plastic cup (see below) and checked for a newly laid egg sac thrice per week. We carefully placed each predominantly first-laid egg sac (for six females we also used the second-laid egg sac due to low survival from the first egg sac) into a 200 ml vial with foam cover, which we sprayed twice a week until hatching. In many species, newly hatched spiderlings remain aggregated before dispersal [59], which appeared crucial for survival in *Nephilingis* [35]. After two weeks of communal rearing (which may have introduced common environmental effects among siblings; see below), were randomly took 20 spiderlings from each full-sib family and transferred them to single-rearing cups, where we monitored each individual five times per week for moults.

For single-rearing of individuals, we used upside down transparent plastic cups (250 ml) with a cotton-filled hole on the top for air and water exchange. Twice a week, we sprayed the cotton with water and fed the spiders. Specifically, all males and female juveniles up to the 4^th^ moult were fed *ad libitum* with *Drosophila sp*., whereas females between the 4^th^ and 6^th^ moult (i.e., two or one moults before reaching maturity; absolute number of moults to maturity vary) were fed blow flies. Females that were one to two moults before maturity were fed two flies, whereas adult females received two or three flies during the first three years of laboratory rearing (see below for thereafter). In the laboratory, we controlled both the temperature (mean = 25 °C, sd = 2 °C) and the light:dark regime (12:12 h). However, some natural light reached the vials (resulting light regime in **Figure 1**).

To test for sex-specific maternal environmental effects, we applied a food treatment during the last three of the total six years of the experiment. In spiders, vitellogenesis occurs predominantly after mating and only in the presence of sufficient food supply [60]. Thus, we mated females within the first three weeks after reaching maturity and subjected them to two maternal food treatments thereafter by feeding them either one (low food) or three flies (high food) twice per week.

### Traits assessed

Between December 2017 and October 2022, we recorded data on adult body mass for 2,540 individuals (789 females, 1,751 males). More data for males were recorded because more males than females survived to adulthood, likely due to the much shorter male developmental time. After reaching sexual maturity, defined by the final moult, somatic growth of both sexes stops but mass may change thereafter (via body condition). We therefore defined adult body size as mass expressed within two days after reaching sexual maturity. We quantified individual adult body size as mass using an analytical balance (KERN ABT-100-5NM; d = 0.00001 g, e = 0.001 g, min = 0.001 g, repeatability = 0.00005 g) located on an anti-vibration table and calibrated before each use.

### Statistical analyses

We were preliminarily interested in the sex-specific relative importance of direct genetic vs. maternal effects on phenotypic variance of adult body size. We were further interested in how strongly these effects are correlated between sexes. To obtain estimates of the required (co)variances, we fitted animal models to adult body size data. The animal model is a mixed model that direct additive genetic or maternal genetic effects (and estimates their variance) via the additive relationships matrix (*A*) and allows simultaneous estimates of fixed effects via generalized least squares solutions [61]. It is possible to statistically separate direct genetic from maternal effects when data exist on related individuals from different mothers [28] and quality of this separation ability depends on size and structure of the pedigree [62]. Further, it is possible to separate maternal environmental from maternal genetic effects and to also estimate the covariance between direct genetic and maternal genetic effects, but data and pedigree requirements increase. In our case, we anticipated to estimate direct genetic and maternal effect variances separately per sex, plus all the possible covariances, thereby increasing data structure requirements, so that we first established what kind of variance model is supported by our data and pedigree structures. We did so by combining approaches of i) model selection among several candidate models, which varied in how we specified the maternal effect variance and whether we included direct-maternal genetic effect covariances, and ii) by data simulations (electronic supplementary material, model selection, data simulations, **Appendix - figures 1-3**).

Using simulations, we were not fully able to separate maternal environmental from maternal genetic effects (electronic supplementary material, **Appendix 1 - figure 3**), and model selection via AIC supported modelling (co)variance of sex-specific maternal effects using one of the simplest approaches considered (electronic supplementary material, model selection, **Table S1**). Specifically, we specified maternal effects as maternal identities, which represent maternal composite effects (i.e., combining putative maternal environmental and maternal genetic effects). In our case, maternal environmental effects may also encompass common environmental effects due to initial common rearing of full-sibs from the same egg sac. We thus modelled sex-specific additive genetic (*a*), maternal (*m*), and residual effects (*e*). For both direct genetic and maternal effects, between-sex covariances can be estimated, whereas this is not possible for the residuals. Accordingly, the assumed multivariate normally distributed random-effect covariance structures for female (F) and male (M) effects with means of zero followed for 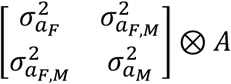, for 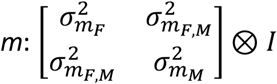, and for 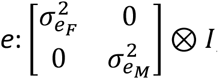, whereby *A* is the pedigree-derived additive relationship matrix and *I* the identity matrix.

We also fitted fixed sex effects and interactions of these sex effects with all other fixed effects that were similar to all candidate models. Specifically, we fitted fixed effects for i) overall sex means (*sex*; female or male), ii) seasonal trends (*date;* integer between 1 and 366, and *sex-by-date*), iii) maternal food treatment (*maternal food*; low or high, and *sex-by-maternal food)*, and iii) experimental period effects (*period*; first or second three-year period, and *sex-by-period*) in respect to the maternal food treatment because the *maternal food* was applied only during the last three of the total six years. We fitted the seasonal trends because development of some spider species, including *Nephilingis*, is affected by day-(or night-) length, i.e., by season [35, 36]. All full siblings hatched on the same day so that season effects may be regarded as either environmentally induced maternal effects or as seasonal common environmental effects, which we wanted to account for here. The *date* trends thus served as general surrogates to many aspects of seasonal day-light variation (**Figure 1**) and enable a more meaningful between-sex comparison by regressing to the common average hatch date.

We modelled adult body mass on the log scale (Ln) because adult body size results from past growth (which may be a proportional process), and the log-scale efficiently accounts for scaling effects both within and between sexes. Within sexes, model residuals based on untransformed data showed a right skew and their variance increased with the fitted values, which also implies variance heterogeneity across seasons (see also raw data in **Figure 1**). Between sexes, the sex-ratio of the untransformed sex-specific standard deviations was of the same magnitude as the ratio of the untransformed sex-specific means (female to male ratio was 56 for standard deviations and 75 for means). The log-transformation stabilized variances both within and between sexes, which accordingly refer to variation in proportional size differences conditional on fitted fixed effects (sex-specific geometric means and systematic trends). Note that an alternatively considered scaling of these mass records, either within or across sexes, does not stabilize variances as does the log transformation. The response vector of natural logarithm of adult body mass (*y*) was modelled as:

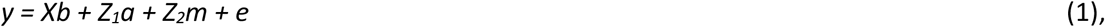

where *X* and *Z* are the design matrices linking data with the fixed and random effects, respectively.

Based on the estimated variance components we calculated the relative contributions per sex (*s*; either female, F, or male, M) of the direct genetic effect variance 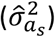 to the total phenotypic variance 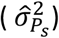, i.e., the heritability (*h*^*2*^), as 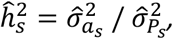 and the corresponding contribution of the maternal effect variance 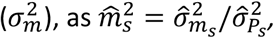 Where 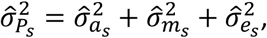 and, 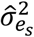is the sex-specific residual variance estimate. We calculated between-sex correlations (*R*_*F,M*_) for genetic 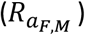 and maternal effects 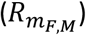 based on the estimates for between-sex covariance 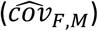 and the sex-specific variances 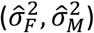, as 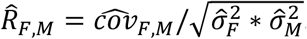. For (co)variance (-based) parameter estimates constrained by boundaries 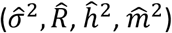, we approximated confidence intervals based on 10,000 parametric bootstrap replicates (electronic supplementary material, parametric bootstraps). We fitted models using residual maximum likelihood (REML) via the average information algorithm implemented in ASReml-R v. 4.1.0.176 [63], executed in R v. 4.1.2, and tested fixed effects using *F*-tests with adjusted denominator degrees of freedom [64].

## Data accessibility

Underlying data and R-scripts (R Markdown file) are available on the Dryad Digital Repository: https://doi.org/10.5061/dryad.tb2rbp039 (during review: https://datadryad.org/stash/share/pXD8qttGbQKGrosfkma4VPY84cdQkIDWr4qeMxCNI). We also provide a html output file of the R Markdown file during review.

## Authors’ contributions

S.K.-F.: funding acquisition, project administration, conceptualization, methodology, supervision, investigation, resources, data curation, writing – original draft, review & editing; M.K.: funding acquisition, resources, writing – review & editing, P.V.D.: methodology, investigation, resources, data curation, formal analysis, visualization, validation, writing – original draft, review & editing.

## Acknowledgements

We thank R. Golobinek, J. Šet, E. Turk, T. Lokovšek, K. Čandek, M. Gregorič and S. Quinones-Lebron for helping with spider collection, rearing, and data recording.

## Appendix 1

### Model selection

We selected the variance structure of the statistical model based on an established model selection criterion (AIC) applied to three candidate models. The models were specified with equal fixed effects (see main text), but different random effects and the complexity of their covariance structures (**Appendix 1 - table 1**). All models estimated sex-specific variance components underlying direct additive genetic (*a*) and residual effects (*e*) as described in the main text. However, sex-specific maternal effects (*m*) we specified as either (or both) maternal environmental effects (*me*), via including a maternal id term, or as maternal genetic effects (*mg*), via a maternal id term and linking ids to the inverse of the pedigree-derived relationship matrix (similarly to what is described in the main text for direct additive genetic effects). We also fitted models with covariances between sex-specific direct genetic and sex-specific maternal genetic effects, which here combines the 2 × 2 covariance matrices for *a* and *m* to a 4 × 4 covariance matrix:

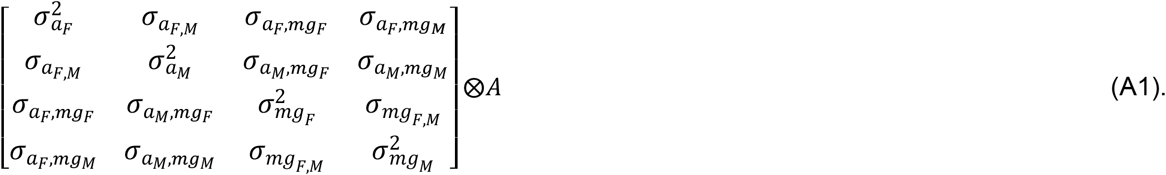

Fitting the covariance(s) between direct genetic and maternal genetic effects may be important because the maternal and direct genetic variance estimates may be biased if the covariance(s), if present, remain unaccounted.

We were unable to consistently fitting models with both maternal environmental and maternal genetic effects. Specifically, results depended on starting values, yielding higher or lower log likelihoods than less complex, nested models, and fitting often aborted because of detected singularities. Thus, we do not report results on models including both types of maternal effects.

The model with maternal environmental effects yielded the lowest AIC (model 1, **Appendix 1 – table 1**), why we reported major results based on this model. Nonetheless, fitting a model with maternal genetic effects and a covariances between direct genetic and maternal genetic effects (model 3) indicated that if maternal effects are at least partly genetic and this covariance is omitted (which was estimated to be negative) additive genetic variance of females may be underestimated considerably (**Appendix 1 - figure 1**) and also female heritability for adult body size (**Appendix 1 - figure 2**). However, estimates based on model 3 also showed high estimate uncertainties for the female direct genetic and residual (co)variance estimates (**Appendix 1 - figure 1**). The strong uncertainty of female relative to male estimates may be explained by fewer records in females than males, and because of a much higher ratio of direct genetic to maternal genetic variance in females, plus an estimated negative covariance between direct genetic and maternal genetic effects – all are expected to lower estimate precision [1].

It should be noted that results based on the alternative models (models 2 & 3) support the conclusion made in the main text about sex differences in adult body size architecture. Specifically, results based on models fitting maternal genetic effects suggested even higher genetic and lower maternal contributions to body size in females and higher maternal and lower genetic contributions in males (**Appendix 1 – figures 1,2**)

### Data simulations

Disentangling maternal environmental from maternal genetic effects requires many data and specific pedigree connections [1-4]. Therefore, we evaluated whether the data and pedigree structures are sufficient to do so. Specifically, we were interested whether we can disentangle between sex-specific maternal genetic effects (*m*_*g*_) and sex-specific maternal environmental (*m*_*e*_) effects. To do so, we used Monte Carlo simulations in which we simulated effects for one variance component at the time: either for additive genetic effects (*a*), maternal genetic effects (*mg*), or maternal environmental effects (*me*). Using the simulated data, we then fitted three models to each data type to estimate, in addition to residual variance, either one variance (the simulated component) or two variance components (the simulated plus one of the two non-simulated components).

For data simulations, we neglected the fixed effects of the model, except for sex-effects, and generated multivariate sex-specific random effects for either *a, mg*, or *me*, and always for residuals (*e*), whereby we drew the sex-specific random effects for *a, mg*, and *me* from a multivariate normal distribution with variance for each sex of 0.25 and a between-sex covariance of 0.125 (i.e., a between-sex correlation of 0.5). Sex-specific residuals were always drawn independently among all individuals from a normal distribution with variance of 0.75. Sex-specific *a* and *mg* were drawn independently among founders and their inheritance was simulated following Mendelian expectations. In detail, all offspring inherit the average of the sex-specific (direct or maternal) genetic effects from both their parents plus a random Mendelian sampling effect drawn from a multivariate normal distribution with half the variance (and half the between-sex covariance) as the founder genetic effects. Thus, the common sex-specific effect each parent passes on to all its sex-specific offspring consist of half of the sum of their average parental sex-specific effects, and each offspring is assigned an own random Mendelian sampling effect, which together make up the (direct or maternal) genetic effect of the offspring. Please note that both males and females inherit sex-specific direct and maternal genetic effects to offspring of both sexes. In contrast, sex-specific *me* were drawn independently among all dams and assigned to their sex-specific offspring. All sons or daughters expressed their own sex-specific direct genetic effect but the sex-specific maternal environmental or maternal genetic effect of their mother, i.e., expressed sex-specific maternal effects were common among brothers or sisters.

Based on model estimates for simulated data, we concluded that our data and pedigree structures are insufficient to reliably disentangle the two maternal effect types. Specifically, simulated maternal environmental effects (*me*) were misinterpreted as (non-simulated) maternal genetic effects (*mg*), but not *vice versa* (**Appendix 1 - figure 3**). Thus, for our data, we must consider that if maternal environmental variance is truly present, maternal genetic variance may be detected statistically even when truly absent.

### Parametric bootstraps

Because confidence intervals for bounded parameters, such as (co)variance-ratio-derived parameters (e.g., correlations, variance proportions) may not be well approximated using frequentists methods (such as the delta method), we obtained confidence intervals for these parameters using parametric simulations [5]. For each of 10,000 parametric bootstrap replicates, we neglected the fixed effects of the model except for sex-effects and generated multivariate random effects for *a, m*, and *e* under the model estimated (co)variances, whereby sex-specific *m* were drawn independently among dams (reflecting maternal environmental effects) and sex-specific *a* were drawn independently among founders. Inheritance of *a* was simulated following Mendelian expectations (as described in **data simulations**). We then refitted the model (as described in the main text; model 1 in Appendix 1 - table 1) to the simulated data, extracted the (co)variance-ratio-derived parameters, and defined the 95% confidence interval for each parameter as the interval between the 0.025^th^ and 0.975^th^ percentiles of the 10,000 estimates.

**Appendix 1 - figure 1.**
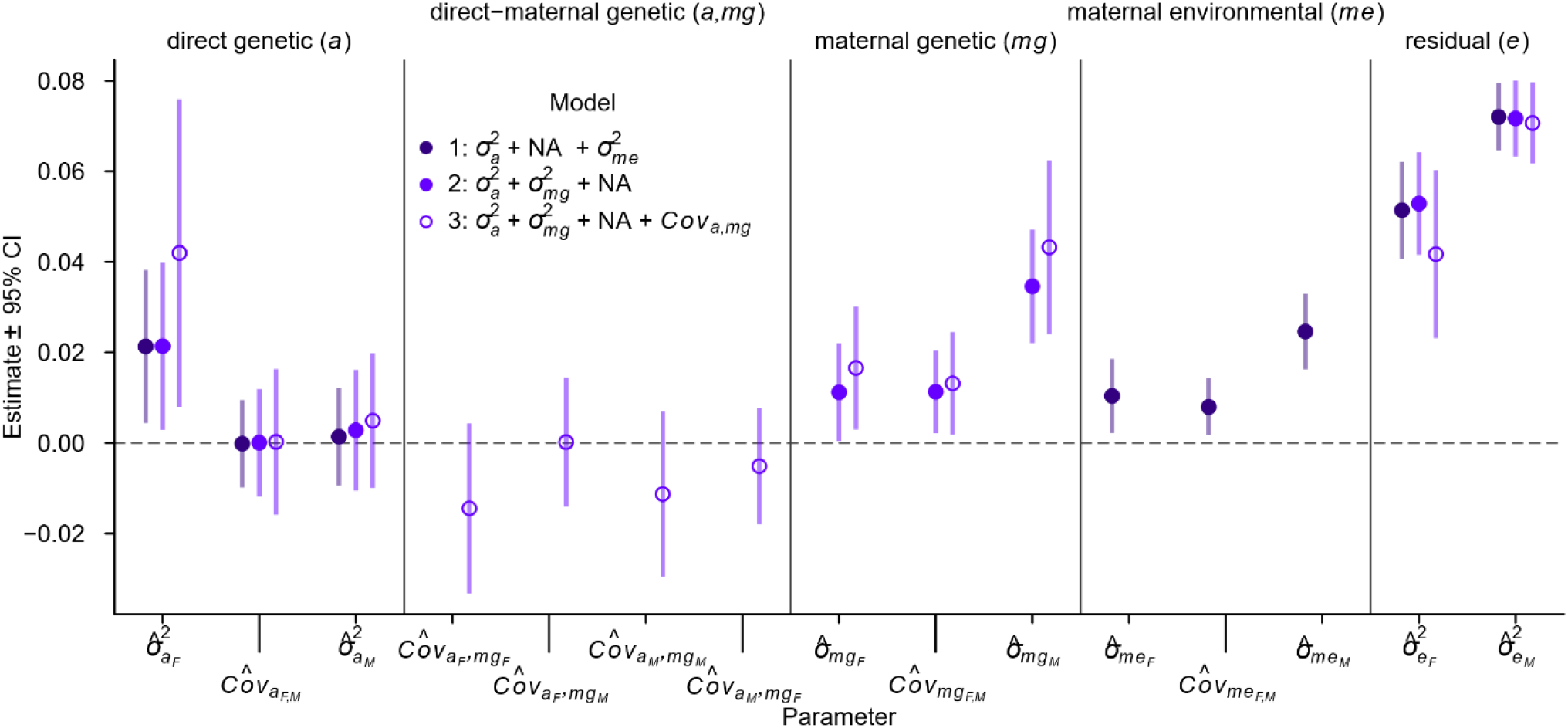
Variance estimates for log of adult body size based on different models. Sex-specific (co)variance estimates based on empirical data using three different models in respect to assuming presence or absence of maternal genetic or maternal environmental effects and covariance between sex-specific direct genetic effects and sex-specific maternal genetic effects. Different colours differentiate between the models as indicated in the legend and results based on model 1 are reported in the results section of the main text. Circles indicate the REML estimates and error bars the delta-method approximate 95% confidence intervals. We acknowledge that the delta method, unlike the parametric bootstrap method, ignores parameter boundaries. Proportional contributions of each variance component to the phenotypic variance and correlations are shown in **Appendix 1 - figure 2**.

**Appendix 1 - figure 2.**
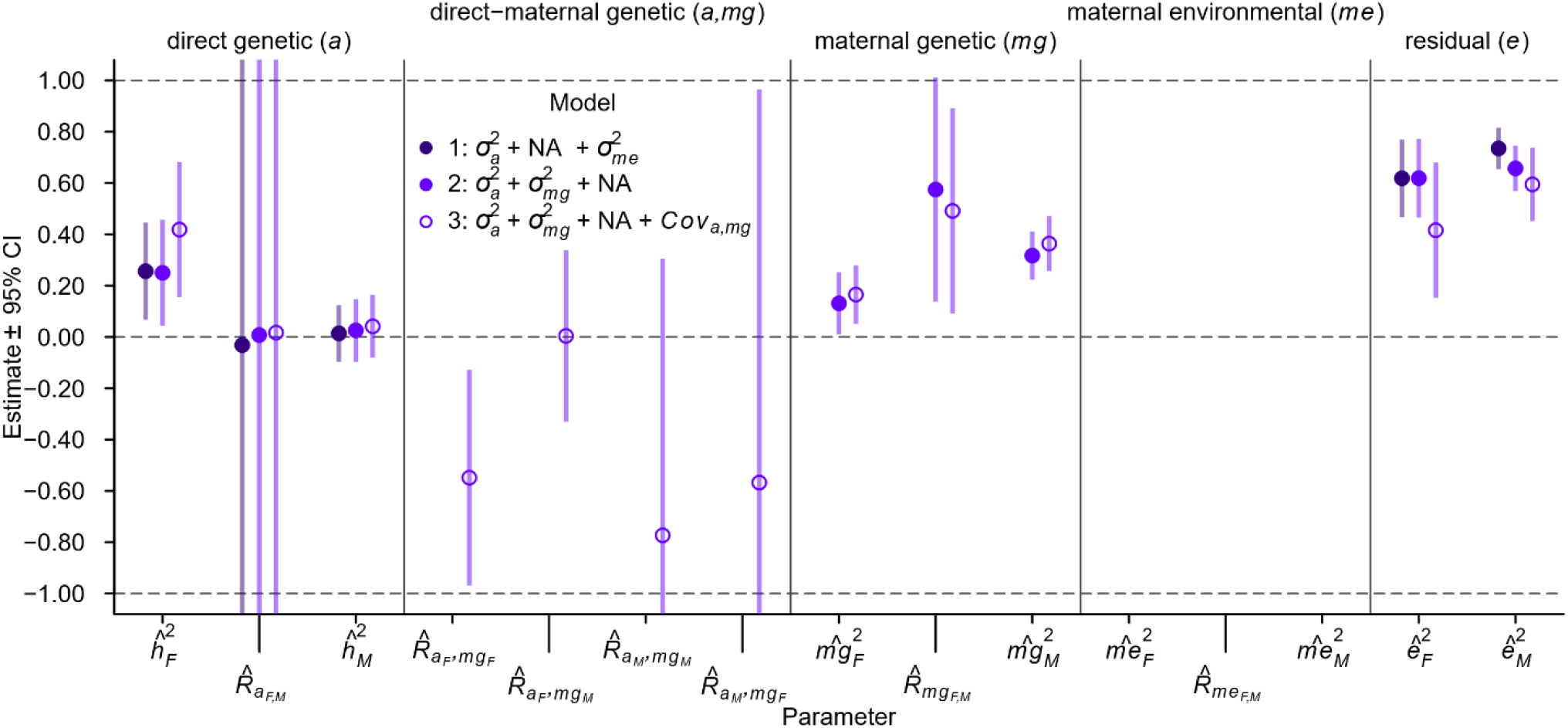
Contribution of variances for log of adult body size based on different models. Sex-specific estimates for proportional contribution to the phenotypic variance and correlations for different random effects based on empirical data using three different models (as in **Appendix 1 - figure 1**) in respect to assuming presence or absence of maternal genetic or maternal environmental effects and covariance between sex-specific direct genetic effects and sex-specific maternal genetic effects. Different colours differentiate between the models as indicated in the legend and results based on model 1 are reported in the results section of the main text. Circles indicate the estimate with approximate 95% confidence intervals according to the delta-method. We acknowledge that the delta method, unlike the parametric bootstrap method, ignores parameter boundaries.

**Appendix 1 - figure 3.**
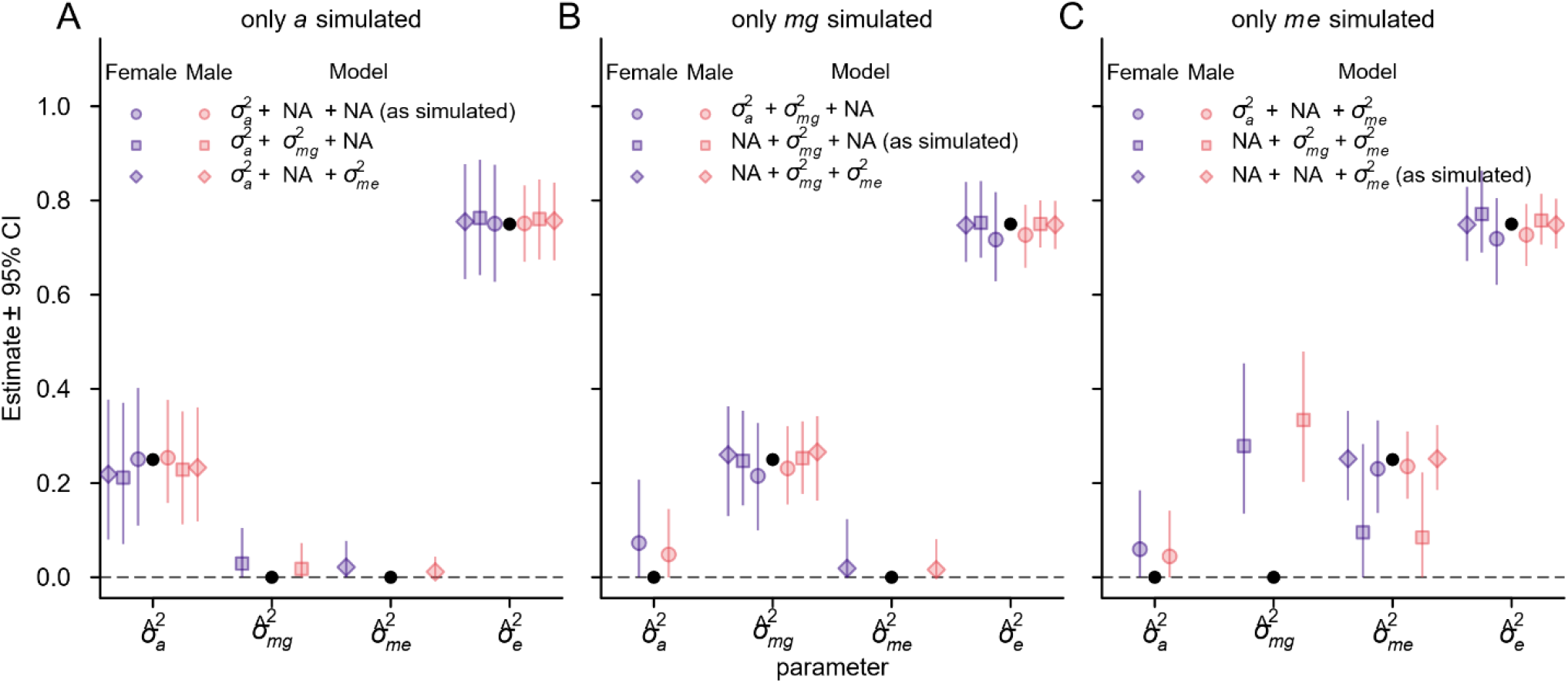
Data simulation results. Sex-specific variance estimates based on simulated data and using either the correct model or incorrect models, each containing one additional non-simulated variance component, to assess estimation bias based on the empirical data structure and pedigree. Simulated were sex-specific effects underlying residual variance 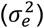 and either direct additive genetic variance 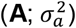 maternal genetic variance 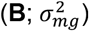, or maternal environmental variance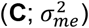. Different symbols differentiate between the models as indicated in the legend (variances were estimated following the covariance matrices as reported in the main manuscript) and simulated variances are indicated by black circles. Error bars show the 95% estimate confidence interval across a maximum of 1,000 simulations (convergence of some simulations using incorrect models failed). The most pronounced bias is present in (**C**), where the 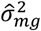 interval is well away from zero, even though only 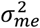 and 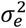 were simulated (i.e., the confidence interval for the estimate 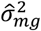 does not cover the true parameter value of 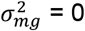).

**Appendix 1 - figure 4.**
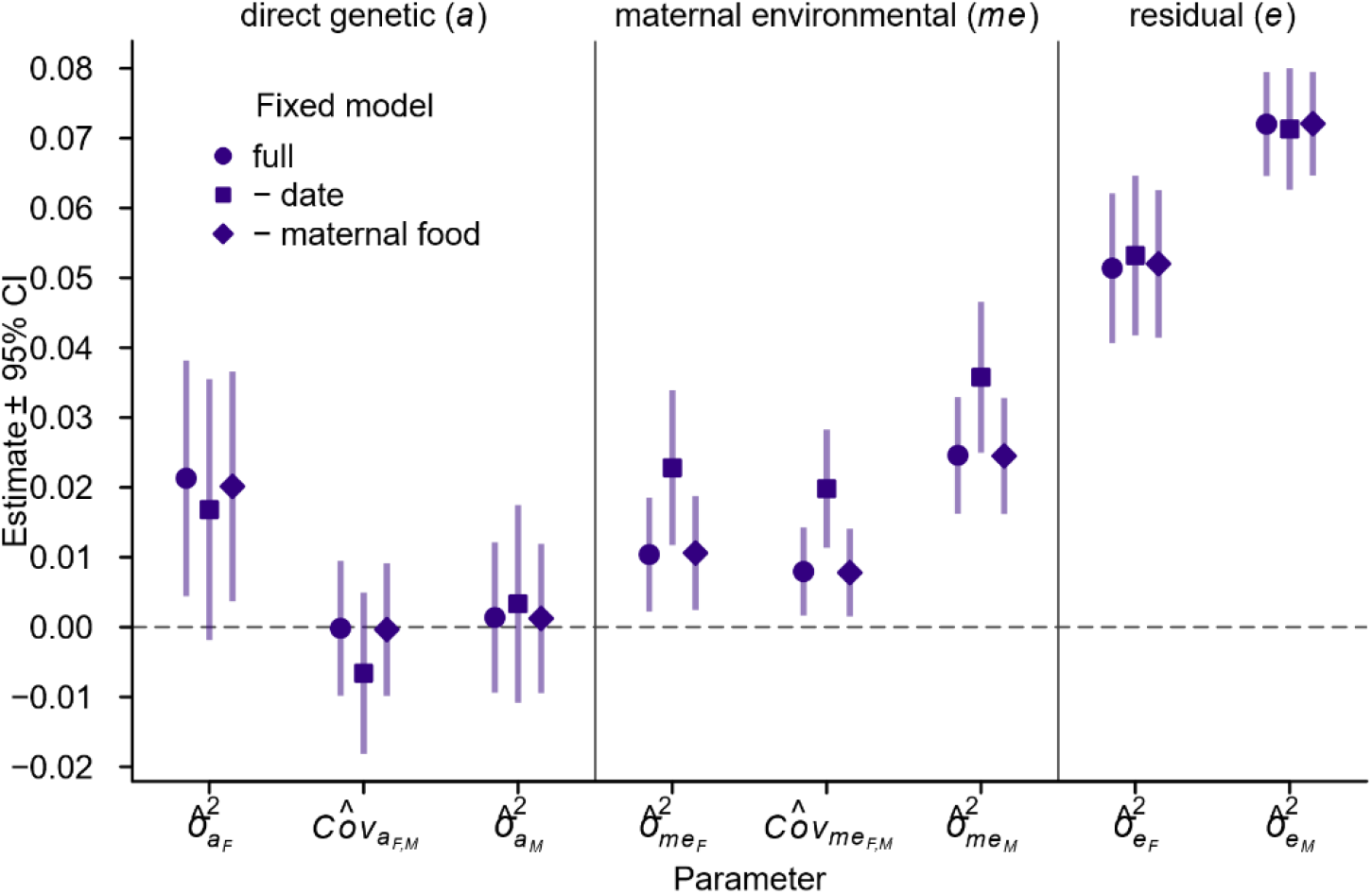
Some variance estimates for log of adult body size depend on included fixed effects. Sex-specific (co)variance estimates based on empirical data for the selected full model (*Full*; model 1 in **Appendix 1 - figure 1**) and when excluding seasonal trends (-date) or maternal food treatment effects (-maternal food). Symbols indicate the REML estimates and error bars the delta-method approximate 95% confidence intervals. We acknowledge that the delta method, unlike the parametric bootstrap method, ignores parameter boundaries.

**Appendix 1 - table 1.** Model-selection parameters for three models on log of adult body size, different in respect to assuming presence or absence of either maternal genetic effects (*mg*) or maternal environmental effects (*me*), and for the latter covariance between sex-specific direct genetic effects (*a*) and sex-specific maternal genetic effects.

| Model | *var(a)* | *var(mg)* | *var(me)* | *cov(a, mg)* | *k* | *logLik* | *AIC* |
| --- | --- | --- | --- | --- | --- | --- | --- |
| 1* | + | - | + | - | 8 | 1876.3 | -3736.5 |
| 2 | + | + | - | - | 8 | 1871.4 | -3726.7 |
| 3 | + | + | - | + | 12 | 1873.6 | -3723.2 |

*var(a)*: sex-specific direct additive genetic effect variance including between-sex covariance. *var(mg)*: sex-specific maternal genetic effect variance including between-sex covariance. *var(me)*: sex-specific maternal environmental effect variance including between-sex covariance. *cov(a, mg)*: covariances between sex-specific direct additive genetic effects and sex-specific maternal genetic effects. *k*: number of (co)variance parameters. *LogLik*: REML estimate of the log likelihood (the higher the better). *AIC*: Akaike's information criterion (the lower the better). \* Chosen model.

